# PURIFICATION METHOD FOR ADENO-ASSOCIATED VIRAL VECTOR SEROTYPE 9 USING CERAMIC HYDROXYAPATITE CHROMATOGRAPHY AND ITS QUALIFICATION

**DOI:** 10.1101/2024.03.12.584728

**Authors:** Yae Kurosawa, Yuji Tsunekawa, Mikako Wada, Yuko Aizen, Yuko Nitahara-Kasahara, Takashi Okada

**Affiliations:** Division of Molecular and Medical Genetics, Center for Gene and Cell Therapy, The Institute of Medical Science, The University of Tokyo, Tokyo 108-8631, Japan; Chromatography Media Business Division, HOYA Technosurgical Corporation, Tokyo 196-0012, Japan

**Author notes:** Corresponding authors: Takashi Okada, Yuji Tsunekawa.

**Keywords:** AAV, purification, TFF, ceramic hydroxyapatite, chromatography

## Abstract

Adeno-associated virus (AAV) vectors can efficiently transduce exogenous genes into various tissues *in vivo*. Owing to their convenience, high efficiency, long-term stable gene expression, and minimal side effects, AAV vectors have become one of the gold standards for investigating gene functions *in vivo*, especially in non-clinical studies. However, challenges persist in efficiently preparing a substantial quantity of high-quality AAV vectors that comprise only genome-packaged full particles. Commercial AAV vectors are typically associated with high costs. Further, in-laboratory production is hindered by the lack of specific laboratory equipment, such as ultracentrifuges. Therefore, a simple, quick, and scalable preparation method for AAV vectors is needed for proof-of-concept experiments. Herein, we present an optimized method for producing and purifying high-quality AAV serotype 9 (AAV9) vectors using standard laboratory equipment and chromatography. Using ceramic hydroxyapatite as a mixed-mode chromatography medium can markedly increase the quality of purified AAV vectors. Basic protocols and optional methods for evaluating purified AAV vectors are also described.

## INTRODUCTION

Adeno-associated virus (AAV), a member of the *Parvoviridae* family, is one of the smallest non-envelope viruses with a single-stranded DNA genome (Samulski & Muzyczka, 2014). AAV has been extensively explored as an *in vivo* gene delivery vector due to its weak immune response, high infective efficiency, and stable long-term gene expression. Consequently, AAV vectors have emerged as promising gene delivery systems for gene therapy modalities (Van Vliet *et al*., 2008). Traditionally, laboratories rely on ultracentrifugation to purify the AAV vectors (Belova *et al*., 2022). Although ultracentrifugation is a sophisticated technique, this process is time consuming, requires expertise, and has low scalability. Although we developed an improved, scalable, and short-term ultracentrifugation method that is compliant with Good Manufacturing Practice (Wada *et al*., 2023), proof-of-concept experiments in the laboratory usually do not require the removal of empty capsids and instead focus on eliminating contaminated proteins and nucleotides. Wada *et al*. (2023) developed a buffer-exchange method using a ceramic hydroxyapatite column as an alternative to dialysis after ultracentrifugation. Thus, in this protocol, we introduced a simple purification method using ceramic hydroxyapatite chromatography (Kurosawa *et al*., 2012; Kurosawa *et al*., 2019). This method enables the separation and purification of AAV vectors at neutral pH, reducing the denaturation and deactivation of AAV vectors that often occur with affinity chromatography (Lins-Austin *et al*., 2020; Miyaoka *et al*., 2023).

## BASIC PROTOCOL 1

### Production of adeno-associated virus serotype 9 vectors in 293EB cells

Previously, we presented our adeno-associated virus serotype 9 (AAV9) production protocol, which outlined the steps for cell culture, triple transfection, and harvesting of the culture supernatant (Wada *et al*., 2023). Using Basic Protocol 1, the culture supernatant can be prepared for the purification process outlined in Basic Protocol 2.

### Materials

HEK293EB cell (Tomono *et al*., 2018)

DMEM (high-glucose) supplemented with L-glutamine and phenol red (Fujifilm Wako Pure Chemical Corporation, cat. no. 044-29765)

Penicillin-streptomycin mixed solution (Nacalai Tesque, Inc., cat. no. 26253-84) Fetal bovine serum (FBS; Sigma-Aldrich Co. LLC, cat. no. F0926-500ML)

Trypsin-ethylenediaminetetraacetic acid (EDTA) (2.5 g/L-Trypsin/1 mmol/L-EDTA solution; Nacalai Tesque, Inc., cat. no. 32777-15)

PBS (D-PBS(−); Nacalai Tesque, Inc., cat. no. 14249-24)

200 mmol/L l-Alanyl-l-glutamine solution (100×) (Nacalai Tesque, Inc., cat. no. 04260-64) NaHCO_3_ (Nacalai Tesque, Inc., cat. no. 31213-15)

D-(+)-glucose (Nacalai Tesque, Inc., cat. no. 16806-25) pHelper (pAdDeltaF6; Addgene, cat. no. 112867) pAAV2/9n (Addgene, cat. no. 112865)

pAAV-ZsGreen1 Vector (Takara Bio Inc., cat. no. 6231)

PEI MAX Transfection Grade Linear Polyethyleneimine Hydrochloride (MW40,000) (Polysciences Inc., cat. no. 24765-1)

Tissue culture flasks (300 cm^2^; T300; TPP Techno Plastic Products AG, cat. no. 90301)

CELLdisc, 4 layers, 1000 cm^2^ (Greiner Bio-One International GmbH, cat. no. 678104)

Countess™ 3FL Automated Cell Counter (Thermo Fisher Scientific Inc., cat. no. AMQAF2000)

Trypan Blue Stain (0.4%) for use with the Countess™ Automated Cell Counter (Thermo Fisher Scientific Inc., cat. no. T10282AMQAF2000)

Countess™ Cell Counting Chamber Slides (Thermo Fisher Scientific Inc., cat. no. C10312)

CO_2_ incubator

500 mL rapid-flow bottle top filter 0.45 μm PES membrane (Thermo Fisher Scientific Inc., cat. no. 295-4545)

Centrifuge (Eppendorf Himac Technologies Co. Ltd.; cat. no. himac CR30NX)

#### Production schedule for AAV9 vectors

Day 1. Preparation of 293EB cells

Day 4. Transfection of the AAV9 vectors into cells Day 9. Harvesting of the AAV9 vector particles

#### Preparation of 293EB cells

1. Seed 293EB cells in culture medium in a CELLdisc (4 × 10^7^ cells/150 mL/CELLdisc) and incubate at 37 °C in a CO_2_ incubator (Day 1).

#### Transfection of the AAV9 vectors into cells

2. On Day 4, transfect three plasmids (pHelper, pAAV2/9n, and pAAV-ZsGreen1) into 293EB cells as described in steps 3–8.
3. Prepare DNA mixture comprising 38.66 μg pHelper, 19.36 μg pAAV2/9n, and 19.36 μg pAAV-ZsGreen1 in 4.4 mL of transfection medium for one CELLdisc.
4. Dilute 116 μL of 2 mg/mL PEI-MAX (total: 232 μg) with 4.4 mL of transfection medium for one CELLdisc.
5. Combine the DNA mixture and diluted PEI-MAX solution, vortex, and allow the mixture to settle for at least 15 min at room temperature.
6. Add the DNA and PEI-MAX mixture to 150 mL of transfection medium in a bottle.
7. Remove the supernatant from the CELLdisc and add 150 mL of transfection medium containing the DNA and PEI-MAX mixture.
8. Culture cells at 37 °C in a CO_2_ incubator.

### Harvesting of the AAV9 vector particles

9. On Day 9, collect and decant the supernatant in a centrifuge bottle.
10. Centrifuge the supernatant at 2,000 × g for 15 min at 4 °C and filter the solution using a 0.45-μm bottle-top filter and a vacuum to remove cells and cell debris.
11. Store the clarified supernatant at 4 °C until use.

## BASIC PROTOCOL 2

### Concentration and buffer exchange of the AAV9 vectors from 293EB cell culture supernatants using tangential flow filtration

A description of the basic protocol for performing the initial step of AAV vector purification is provided below. A tangential flow filtration (TFF) system is used to concentrate and diafiltrate the AAV9 crude sample. This process yields the AAV9 vector solution required for the subsequent step: Basic Protocol 3.

### Materials

HEPES (Dojindo Laboratories, cat. no. 346-08235)

MES (Nacalai Tesque, Inc. cat. no. 21623-26)

1 M MgCl_2_ (Nacalai Tesque, Inc. cat. no. 20942-34)

NaCl (Nacalai Tesque, Inc. cat. no. 31320-05)

NaOH (Nacalai Tesque, Inc. cat. no. 31511-05)

Ethanol (EtOH; Nacalai Tesque, Inc.cat. no. 14813-95)

Water bath (Thermax Water Bath TM-2A, AS ONE Corporation, cat. no. 1-4594-32)

ReadyFilter Hollow Fiber Cartridge 500 K, membrane surface area of 650 cm^2^ (Cytiva, cat. no. RTPUFP-500-C-4MS)

KrosFlo^®^ KR2i TFF System (Repligen Corporation)

500-mL rapid-flow bottle top filter with 0.45-μm PES membrane (Thermo Fisher Scientific Inc., cat. no. 295-4545)

#### Concentration and buffer exchange of the AAV9 vector from 293EB cell culture supernatants using tangential flow filtration

A solution comprising 10 mM HEPES, 150 mM NaCl pH 7.2, and 1 mM MgCl_2_ or 10 mM MES, 150 mM NaCl pH 6.5, and 1 mM MgCl_2_ can be used as the TFF buffer.

1. Pre-warm 500 mL of clarified cell culture supernatant from Basic Protocol 1 and TFF buffer (10 mM MES, 150 mM NaCl pH 6.5, and 1 mM MgCl_2_) at 37 °C.
2. Connect the hollow fiber cartridge (500K) to the TFF system.
3. Rinse the hollow fiber cartridge with ultrapure water at a flow rate of 200 mL/min to remove 30% EtOH.
4. Concentrate approximately 500 mL of the cell culture supernatant to 50–60 mL at a flow rate of 800 mL/min.
5. Diafiltrate the concentrated culture supernatant with 2 L of pre-warmed TFF buffer at a flow rate of 800 mL/min.
6. Collect the diafiltrated sample and add 50 mL of the fresh TFF buffer to the sampling bottle.
7. Circulate the buffer at a flow rate of 200 mL/min for 10 min by closing the permeate vent to collect the remaining AAV vector in the system.
8. Repeat steps 6 and 7.
9. Collect the remaining AAV vector in the system.
10. Filter the diafiltrated culture supernatant (total: 200 mL) through a 0.45-μm bottle top filter.
11. After TFF, store the sample at 4 °C until use.

*After TFF, 0*.*5 M NaOH pre-warmed at 55 °C should be pumped into the system and lines at a flow rate of 200 mL/min for 30–60 min, followed by ultrapure water at a flow rate of 200 mL/min for 30–60 min and finally 30% EtOH. The cartridge can then be stored at 4 °C*.

## BASIC PROTOCOL 3

### Purification of the AAV9 vectors from TFF samples using ceramic hydroxyapatite chromatography

The basic protocol used to perform the second step of AAV vector purification is outlined below. Ceramic hydroxyapatite chromatography is employed to purify AAV9 vectors from the concentrated and diafiltrated crude supernatants using TFF. This step yields AAV9 vectors with high purity. The evaluation protocol is described in Basic Protocol 4.

### Materials

CHT™ Ceramic Hydroxyapatite, Type I, 40 μm (Bio-Rad Laboratories, Inc.)

Na_2_HPO_4_·12H_2_O (Nacalai Tesque, Inc., cat. no. 31722-45)

NaH_2_PO_4_·2H_2_O (Nacalai Tesque, Inc., cat. no. 31718-15)

NaCl (Nacalai Tesque, Inc., cat. no. 31320-05)

HEPES (Dojindo Laboratories, cat. no. 346-08235)

NaOH (Nacalai Tesque, Inc., cat. no. 31511-05)

EtOH (Nacalai Tesque, Inc. cat. no. 14813-95)

Empty stainless-steel column (4.6 mm i.d. × 35 mm, 10 μm mesh filter; Sugiyama Shoji Co., Ltd.) Vantage L Laboratory Column (VL16 × 250; Merck KGaA)

AKTA avant 25 (Cytiva)

#### Packing of the empty column with ceramic hydroxyapatite media

1. To create a small ceramic hydroxyapatite column, pack CHT media into an empty stainless-steel column (in-house) using a dry method.

*To create a large column, pack CHT media into a Vantage L Laboratory Column (in-house) using a wet method*.

#### Setting up the chromatography system; AKTA avant 25

2.Place each pump and buffer line in the chromatography system in the appropriate buffer. Equilibrate the sample loop, pH probe, and fraction collector line with 10 mM HEPES, 150 mM NaCl pH 7.2 (equilibration buffer).

#### Purification of the AAV9 vectors using ceramic hydroxyapatite chromatography

If the buffer comprising 10 mM MES, 150 mM NaCl pH 6.5, and 1 mM MgCl_2_ is not used for the TFF step, the pH of the sample should be adjusted to approximately 6.5 after TFF.

1. Place a small ceramic hydroxyapatite column in AKTA avant 25, wash the column with 5 mL of 0.5 M NaOH (CIP solution) and 10 mL of 400 mM sodium phosphate buffer (NaP) pH 7.2 (wash buffer), and equilibrate with the equilibration buffer (> 15 mL) at a flow rate of 1.0 mL/min.
2. Load 10 mL of the sample onto the small ceramic hydroxyapatite column.
3. Wash the column with 10 mL of the equilibration buffer.
4. Elute the AAV9 vector using 10 mL of 15 mM NaP and 150 mM NaCl pH 7.2 (15% B pump of elution buffer (100 mM NaP, 150 mM NaCl pH 7.2)).
5. Wash the column with 5 mL of wash buffer.

Chromatograms are presented in Figure 1. Basic Protocol 4 can be used to determine the titer of the AAV9 vector in each fraction.

**Figure 1.**
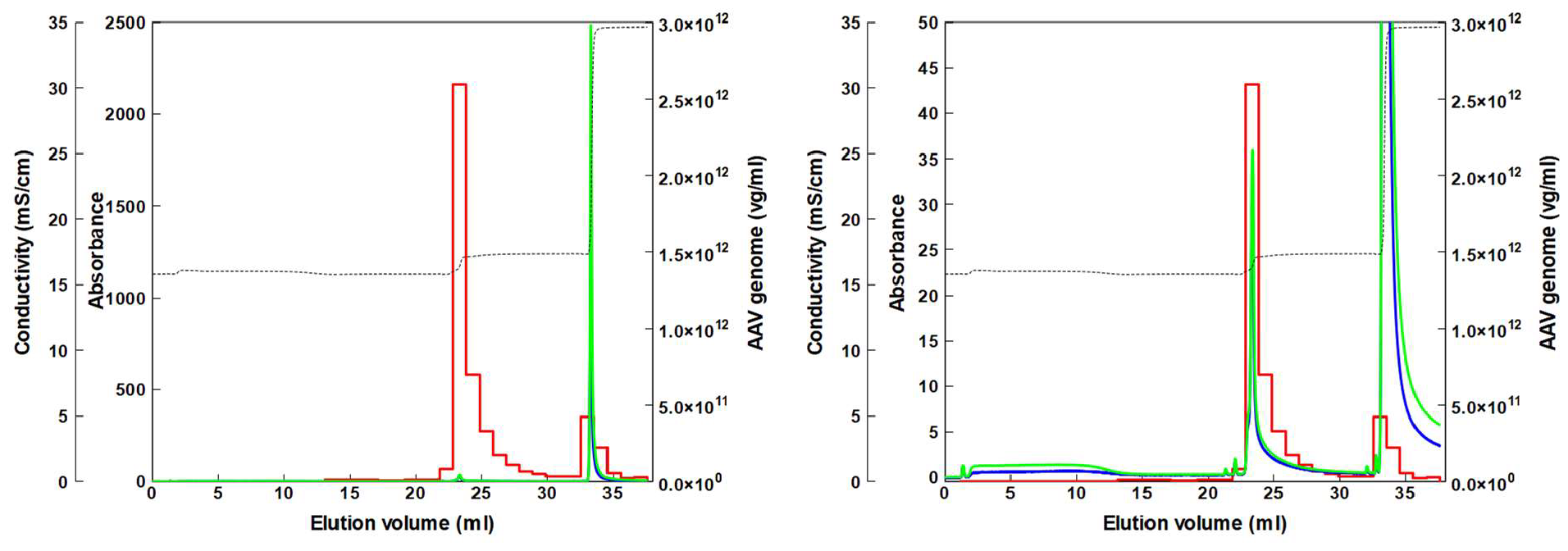
Chromatogram of the AAV9 vectors after TFF on a ceramic hydroxyapatite column. The TFF sample (10 mL) was loaded onto a ceramic hydroxyapatite column (4.6 mm i.d. × 35 mm) washed with 10 mL of 10 mM HEPES, 150 mM NaCl pH 7.2 (10 mL), eluted with 15 mM NaP and 150 mM NaCl pH 7.2 (10 mL), and washed with 400 mM NaP pH 7.2 (5 mL). For each of the fraction obtained the titer of the AAV9 vector was quantified using qPCR (Basic Protocol 4). Red line, AAV9 titer based on qPCR; blue line, ultraviolet (UV) absorbance at 280 nm; green line, UV at 260 nm; black broken line, conductivity. Left: overall; right: magnified UV axis

*When a large ceramic hydroxyapatite column is used, the buffer and sample volumes should be increased by 12-fold*.

*CIP solution should be pumped into the column and system. To remove the alkaline solution, the column and system should be washed with ultrapure water (0.22 μm-filtered and degassed) after chromatography. Finally, 20% EtOH should be injected*.

## BASIC PROTOCOL 4

### Quantification and evaluation of the purified AAV9 vector

The basic protocols for quantifying and evaluating the purified AAV vectors are outlined below. The copy number of the viral genome (vg) for the purified AAV vector can be quantified using TaqMan quantitative polymerase chain reaction (qPCR). The amount of protein contaminants in the AAV vector solution can be determined using the HEK293 HCP ELISA Kit and sodium dodecyl sulfate-polyacrylamide gel electrophoresis (SDS-PAGE), according to the manufacturer’s instructions. The amount of double-stranded DNA (dsDNA) contaminants derived from the host cell dsDNA and plasmid DNA can be quantified using a Quant-iT PicoGreen dsDNA Assay Kit. Finally, the full/empty ratio of purified AAV9 vectors can be determined using transmission electron microscopy (TEM) (Wada *et al*., 2023).

### Materials

Recombinant DNase I (RNase-free) (Takara Bio Inc.; cat. no. 2270A)

DEPC-treated water (Nacalai Tesque Inc., cat. no. 36415-54)

Buffer AL lysis buffer (QIAGEN, cat. no. 19075)

THUNDERBIRD probe qPCR Mix (TOYOBO Co., Ltd., cat. no. QPS-101)

Detection probe for the ZsGreen1 transgene: 5’-/56-FAM/TT CAT CCA G/ZEN/C ACA AGC TGA C/3IABkFQ/-3’ (Bryda *et al*., 2019) (synthesized by Integrated DNA Technologies, Inc.)

Primers for the ZsGreen1 transgene: 5’-GTG TAC AAG GCC AAG TCC GT-3’ and 5’-CCA CTT CTG GTT CTT GGC GT-3’ (Bryda *et al*., 2019) (synthesized by Eurofin Genomics K.K.)

NuPAGE LDS Sample buffer (4×) (Thermo Fisher Scientific Inc., cat. no. NP0007)

NuPAGE Reducing Agent (10×) (Thermo Fisher Scientific Inc., cat. no. NP0009)

NuPAGE MOPS SDS Running Buffer (20×) (Novex, cat. no. NP0001)

NuPAGE 4-12% Bis-Tris Gel 1.5 mm, Mini Protein Gel, 10-well (Thermo Fisher Scientific Inc., cat. no. NP0335BOX)

PageRuler Unstained Protein Ladder (Thermo Fisher Scientific Inc., cat. no. 26614)

Oriole fluorescent gel stain (Bio-Rad Laboratories, Inc., cat. no. 1610496)

Quant-iT PicoGreen dsDNA Assay Kit (Thermo Fisher Scientific Inc., cat. no. P7589)

Black-type fluorescent plate S (Sumitomo Bakelite Co., Ltd., cat. no. MS-8496K)

HEK293 HCP ELISA Kit 3G (Cygnus Technologies, LLC, cat. no. F650S)

Sample Diluent Buffer (Cygnus Technologies, LLC, cat. no. I700)

Phosphotungstic Acid EM (TAAB Laboratories Equipment Ltd., cat. no. P013)

MicroAmp Fast Optical 96-Well Reaction Plate, 0.1 mL (Thermo Fisher Scientific Inc., cat. no. 4346907)

MicroAmp Optical Adhesive Film (Thermo Fisher Scientific Inc., cat. no. 4311971)

Thermal Cycler (C1000 Touch; Bio-Rad Laboratories, Inc.)

QuantStudio 3 (Applied Biosystems by Thermo Fisher Scientific Inc.)

Heat block (block incubator; Astec Co., Ltd., cat. no. BI-516S)

Electrophoresis system (XCell SuperLock Mini-Cell; Thermo Fisher Scientific Inc., cat. no. EI0001)

PowerStation Ghibli I (ATTO Corporation, cat. no. WSE-3100)

ChemiDoc™ Touch Imaging System (Bio-Rad Laboratories, Inc., cat. no.1708370)

Plate reader (Varioskan Flash; Thermo Fisher Scientific Inc.)

Collodion membranes (Nissin EM Co., cat. no. 6512)

Ion bombarder (Nissin EM Co., type PIB-10)

Filter paper (Advantec Toyo Kaisha, Ltd., cat. no. 00011090)

TEM (Hitachi High-Tech Corporation, cat. no. HT7800)

#### qPCR

1. Add recombinant DNase I (RNase-free; 1 μL), 10 × DNase I buffer (2 μL) and DEPC-treated water (15 μL) to the samples (2 μL each) and incubate the mixture at 37 °C for 15 min and 95 °C for 10 min in a thermal cycler.
2. Lyse the vector particles (DNase-treated sample 2 μL) with AL Lysis buffer (40 μL) and DEPC-treated water (38 μL) at 56 °C for 10 min in a thermal cycler.
3. Dilute the lysed sample 100-fold using DEPC-treated water.
4. Prepare the pre-mixed reagents in Table 1.
5. Dispense 18 μL of the pre-mixed reagents and 2 μL of the diluted samples into the wells of a 96-well microplate.
6. Place the sealed microplate in the QuantStudio 3 instrument, perform PCR according to the conditions outlined in Table 2 (recommended by TOYOBO Co., Ltd.), and determine the titers.

**Table 1.** Pre-mixed reagents for qPCR.

| Reagents | Vol (μL) |
| --- | --- |
| TOYOBO Thunderbird probe qPCR mix (2×) | 10 |
| ZsGreen1-probe (10 μM) | 0.4 |
| ZsGreen1-F primer (100 μM) | 0.06 |
| ZsGreen1-R primer (100 μM) | 0.06 |
| ROX reference dye (50×) | 0.04 |
| DEPC-treated water | 7.44 |
| Total | 18 |

**Table 2.** PCR cycling conditions.

| Step |  | Temperature (°C) | Time (s) |
| --- | --- | --- | --- |
| Initial denaturation |  | 94 | 20 |
| PCR<br>(40 cycles) | Denaturation | 95 | 3 |
|  | Extension | 60 | 30 |

#### SDS-PAGE

*Perform the following steps using 0.5–5.0 × 10*^*10*^ *vg/lane of samples:*

1. Preheat heat block to 70 °C.
2. Prepare the sample buffer mixture by mixing 8.75 μL of NuPAGE LDS Sample Buffer (4×) and 3.5 μL of NuPAGE Reducing Agent (10×) per sample.
3. Dispense 12.25 μL of the sample buffer mixture into 1.5 ml microtubes.
4. Dispense 22.75 μL of the sample into microtubes.
5. Heat the mixture at 70 °C for 10 min on the heat block.
6. Dilute the NuPAGE MOPS SDS Running Buffer 20-fold to prepare the running buffer.
7. Assemble the electrophoresis system by adding the gel to the system and pouring the running buffer into the tank.
8. Dispense 2 μL/lane of marker (ladder) and 35 μL/lane of samples into the appropriate wells.
9. Place the lid on the system, connect the cords to the power station, and run electrophoresis at 200 V (constant voltage) for 50 min.
10. Remove the gel and gel plates from the system.
11. Stain the gel using an oriole fluorescent gel stain with shaking for 90 min at room temperature.
12. Lightly rinse the gel with ultrapure water.
13. Place the gel in the ChemiDoc™ Touch Imaging System and capture an image.

*Analyze the purity of the product using SDS-PAGE, as shown in Figure 2*.

**Figure 2.**
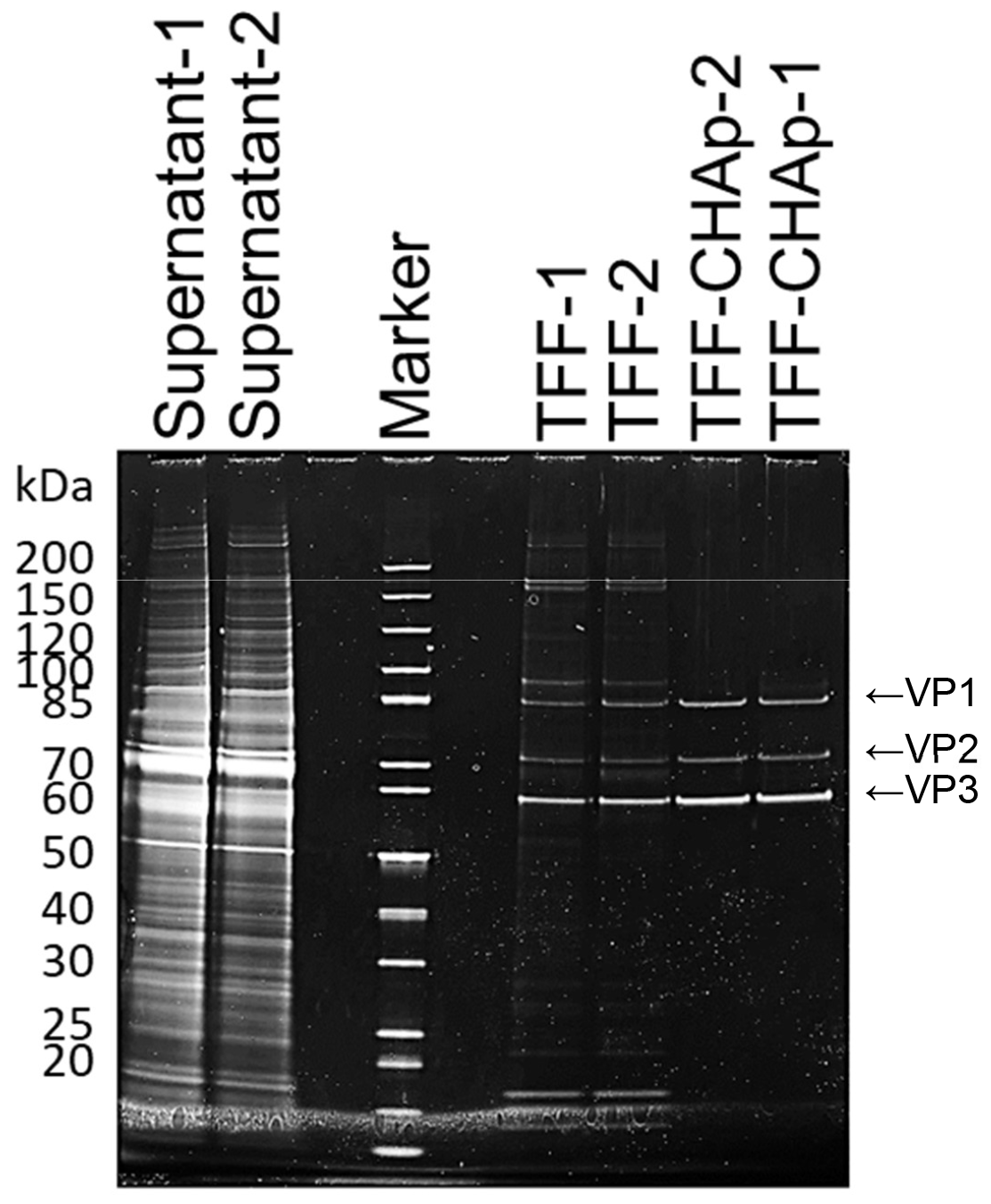
SDS-PAGE analysis of each purification step. The protein contaminants from each purification step were visualized via Oriole staining. SDS-PAGE was performed using a 4–12% gel; 35 *μ*L of sample was loaded into the appropriate lane. The samples were then stained with an Oriole fluorescent gel stain for 90 min. TFF, sample after TFF; TFF-CHAp, pooled fractions from ceramic hydroxyapatite chromatography. The numbers indicate the number of experiments. Arrows indicate the AAV structural proteins: VP1, VP2, and VP3.

#### Quantifying the level of the host cell protein contaminant

*Use the HEK293 HCP ELISA Kit according to the manufacturer’s instructions, with minor modifications*.

1. Allow all reagents to equilibrate to room temperature.
2. Prepare the wash buffer (1 L) by diluting 50 mL of the wash concentrate (20×) with ultrapure water.
3. Dispense 100 μL of anti-HEK:HRP into each well.
4. Dispense 50 μL of the kit standards and samples into the appropriate wells and incubate the microplate at room temperature in the dark for 1.5 h.
5. Wash the plate four times with 200 μL/well of wash buffer.
6. Dispense 100 μL of the TMB substrate into each well and incubate the microplate at room temperature for 30 min.
7. Stop the reaction to by adding 100 μL of stop solution to each well.
8. Measure the absorbance at 450/650 nm using a microplate reader.
9. Calculate the concentration of host cell protein (HCP) in each sample using the standard curve.

#### Quantifying the level of the dsDNA contaminant

*Use the Quant-iT PicoGreen dsDNA Assay Kit according to the manufacturer’s instructions*.

1. Dilute the TE supplied in the kit by 20-fold with ultrapure water (1 mL TE+19 mL ultrapure water, per plate).
2. Dilute the DNA standard curve for calibration with TE to a final volume of 100 μL in microplate wells.
3. Dilute the samples with TE to a final volume of 100 μL in microplate wells.
4. Add 100 μL of PicoGreen diluted 200-fold to each sample.
5. Incubate the samples at room temperature in the dark for 5 min.
6. Measure the fluorescence of the samples using a microplate reader (excitation ∼480 nm, emission ∼520 nm).
7. Determine the concentration of dsDNA in the samples using a standard curve.

*Determine purity by employing commercial kits to confirm the removal of dsDNA and HCP contaminants (Table 3)*.

**Table 3.** Removal of contaminants.

|  | dsDNA |  |  | HCP |  |  |
| --- | --- | --- | --- | --- | --- | --- |
|  | Conc.<br>(ng/mL) | Removal<br>(%) | Total<br>removal<br>(%) | Conc. (ng/mL) | Removal<br>(%) | Total<br>removal<br>(%) |
| Supernatant | 7,309 |  |  | 36,755 |  |  |
| TFF | 1,446 | 92.085 |  | 21.6 | 99.976 |  |
| Chromatography | 3.0 | 99.959 | 99.997 | 8.0 | 92.613 | 99.998 |

#### Full/empty ratio analysis of the AAV9 vectors using transmission electron microscopy (optional)

*Perform transmission electron microscopy (TEM), according to previously described method (Wada et al*., *2023)*.

1. Hydrophilize the collodion membranes using an ion bombarder.xs
2. Place the samples (3 μL) on a hydrophilized grid (collodion membranes) for 1 min.
3. Dispense approximately 3 μL of ultrapure water and adsorb using a filter paper; repeat three times.
4. Stain the samples with 3 μL of 2% phosphotungstic acid solution pH 7.0 for 10 s and then adsorb with a filter paper.
5. Place the membrane on its face and place the sample on a filter paper for 2 min.
6. Analyze the samples loaded onto the collodion membranes using TEM (HC-1 mode; high resolution; acceleration voltage, 100 kV; magnification, 25.0 k).
7. Analyze the full/empty ratio of AAV vectors by counting the full and empty capsid numbers.

*Analyze the full/empty ratio of the purified AAV9 vector using TEM, as shown in Figure 3*.

**Figure 3.**
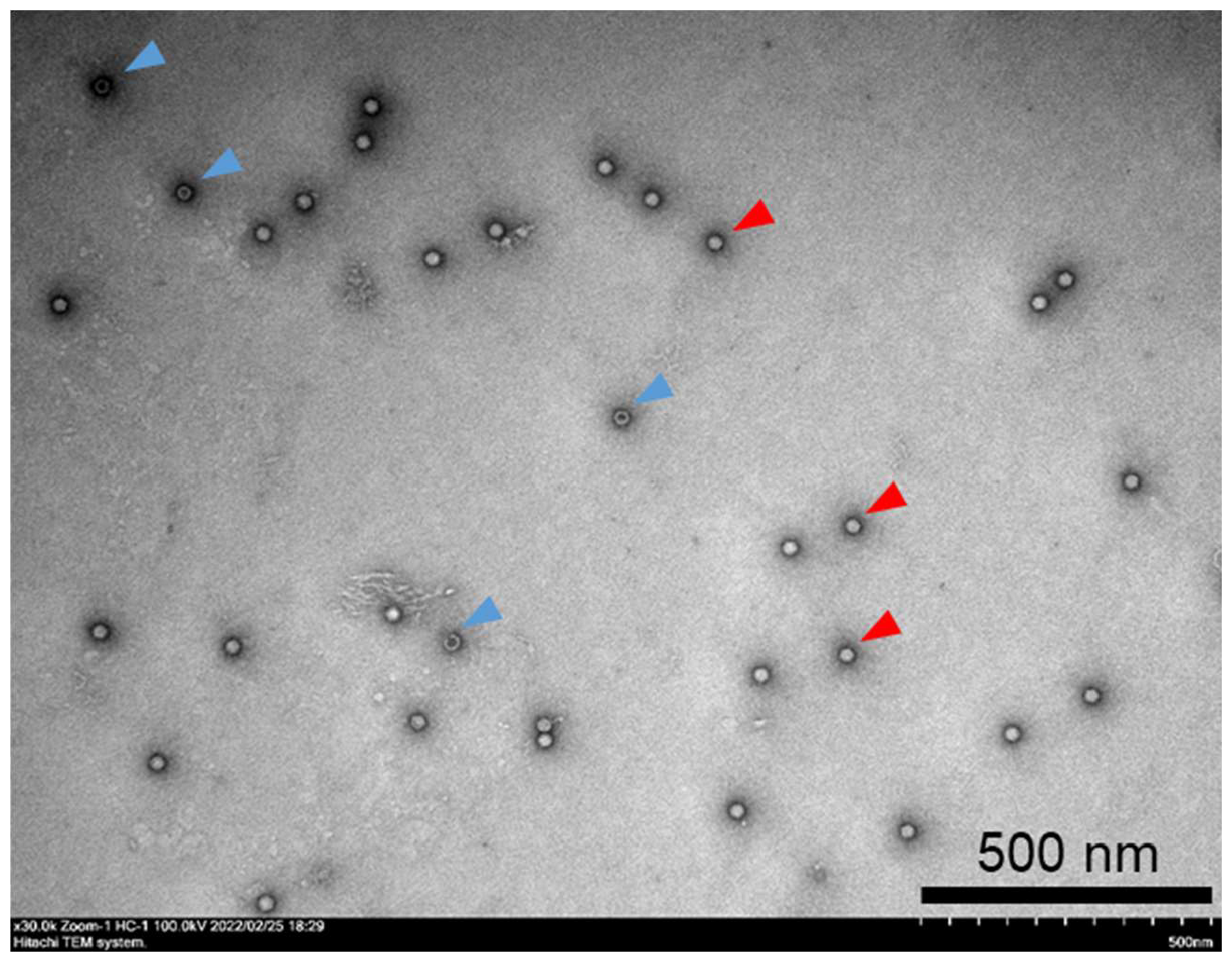
Morphology of the purified AAV9 vector particles based on TEM. AAV9 vectors were loaded onto hydrophilized grids, stained with 2% phosphotungstic acid solution, and observed using TEM at a magnification of 30,000 ×. Red and blue arrowheads indicate full and empty particles, respectively.

## REAGENTS AND SOLUTIONS

### Culture medium

500 mL D-MEM

50 mL FBS

5 mL Penicillin-streptomycin mixed solution Prepare under sterile conditions

### Transfection medium

500 mL D-MEM

5 mL Penicillin-streptomycin mixed solution

5 mL 2 mmol/L-alanyl-L-glutamine solution

8.4 mL 7.5% NaHCO_3_ (filtered with 0.22 μm filter)

6.7 mL 10% D-(+)-glucose (filtered with 0.22 μm filter)

Prepare under sterile conditions

### Stock solution of 2 mg/mL PEI-MAX

1 g PEI MAX Transfection Grade Linear Polyethylenimine Hydrochloride

500 mL Ultrapure water

Adjust the pH to approximately 7.0 and filter the solution (0.22 μm).

Store aliquots at −30 °C.

### 10× TFF stock solution: 100 mM MES and 1.5 M NaCl pH 6.5

21.33 g MES

87.66 g NaCl

800 mL Ultrapure water

Adjust the pH to 6.5 and the volume to 1 L.

### TFF buffer: 10 mM MES, 150 mM NaCl pH 6.5, and 1 mM MgCl2

Dilute the 10× TFF stock solution 10-fold with ultrapure water and add a 1/1,000 volume of 1 M MgCl_2_ (final 1 mM) to the TFF buffer.

*Of note, 10 mM HEPES, 150 mM NaCl pH 7.2, and 1 mM MgCl*_*2*_ *can also be used as the TFF buffer. However, the pH of the sample must be adjusted to 6.5 before loading onto a ceramic hydroxyapatite column*.

### Equilibration buffer: 10 mM HEPES and 150 mM NaCl pH 7.2

4.77 g HEPES

17.53 g NaCl

1,900 mL Ultrapure water

Adjust the pH to 7.2 and the volume to 2 L Filter (0.22 μm) and degas for chromatography

### Elution buffer: 100 mM NaP and 150 mM NaCl pH 7.2

Adjust the pH to 7.2 by mixing 100 mM Na_2_HPO_4_·12H_2_O-150 mM NaCl and 100 mM NaH_2_PO_4_·2H_2_O-150 mM NaCl

Filter (0.22 μm) and degas

### Wash buffer: 400 mM NaP pH 7.2

Adjust the pH to 7.2 by mixing 400 mM Na_2_HPO_4_·12H_2_O and 400 mM NaH_2_PO_4_·2H_2_O. Filter (0.22 μm) and degas

### CIP solution: 0.5 M NaOH

20 g NaOH

Adjust the volume to 1 L using ultrapure water.

Filter (0.22 μm) and degas

## COMMENTARY

### Critical Parameters

Using the correct amount of plasmids and PEI is critical in the production of AAV9 vectors (Basic Protocol 1) as different ratios or amounts can affect the cytotoxicity and efficiency of production.

TFF is an effective tool for removing contaminants, such as host cell proteins (HCPs), DNA, and plasmids, and concentrating the AAV vector in the culture supernatant (Basic Protocol 2). Crucial considerations include pre-warming the buffer and sample at 37 °C. This step increases the fluidity of the sample and buffer and efficiently removes small molecules. A TFF with a 500-kDa cut-off hollow fiber is recommended in this protocol. Performing TFF with a smaller cut-off will increase recovery under the same conditions but will decrease the efficiency of small molecule removal. As this protocol is optimized for a 500 mL sample and a hollow fiber with a membrane surface area of 650 cm^2^, washing with 2 L of buffer should be performed to effectively remove small molecules.

The critical elements in Basic Protocol 3 include the NaP buffer and sample pH for ceramic hydroxyapatite chromatography. Anhydrous sodium dihydrogen phosphate or anhydrous disodium hydrogen phosphate should not be used to prepare NaP buffer due to the potential presence of polyphosphoric acid. This acid can chelate Ca in the apatite and damage the column. The pH of the samples should be adjusted to approximately 6.5. However, replacing the buffer (pH 6.5) with TFF buffer eliminates the need for pH control. A high sample pH decreases the adsorption efficiency while a low sample pH (below pH 6.5) damages the column. To avoid damaging the column, the addition of EDTA to the buffer is discouraged. TEM analysis enables the differentiation between full and empty capsids with small sample volumes (Basic Protocol 4). Tweezers can be used to grasp the membrane at the edge of the grid. When the sample is placed on the membrane, subsequent steps proceed smoothly. An AAV vector concentration of more than 1.0 × 10^12^ vg/mL is ideal for TEM with negative staining. The prepared samples can be stored in a desiccator for an extended duration.

### Troubleshooting Table

The troubleshooting procedure and corresponding solutions are presented in Table 4.

**Table 4.** Troubleshooting.

| Problem | Possible cause | Solution |
| --- | --- | --- |
| **Basic Protocol 2** |  |  |
| Increased pressure in TFF | Membrane is clogged | Wash hollow fiber well with pre-warmed NaOH at 55 °C and rinse with ultrapure water. Follow the manufacturer’s manual. |
| **Basic Protocol 3** |  |  |
| Low adsorption efficiency on the column | Sample pH is not suitable | Check the sample pH and re-adjust to 6.5. |
|  | Excessive loading volume | Decrease and optimize the volume of sample. |
| **Basic Protocol 4** |  |  |
| AAV capsids not found | Low concentration of AAV | Concentrate samples via ultrafiltration (e.g., ultrafiltration column) |

### Understanding the Results

AAV9 vectors with a high titer were produced using Basic Protocol 1 and then purified using Basic Protocols 2 and 3. Figure 1 shows a representative chromatogram of the AAV9 vector after TFF using a ceramic hydroxyapatite column. The eluates were pooled and analyzed via SDS-PAGE (Figure 2). The contaminant bands in the sample decreased after TFF, but remained detectable. Following two-step purification with TFF and ceramic hydroxyapatite chromatography, good purity was obtained and only three AAV structural proteins: VP1, VP2, and VP3, were detected. The purity was further evaluated (Table 3). In terms of contaminants, HCP was efficiently removed by TFF while dsDNA was efficiently removed by ceramic hydroxyapatite chromatography, with removal efficiencies of 99.998% and 99.997%, respectively (Table 3). Morphological analysis of the purified AAV9 vector particles was performed using TEM (Figure 3). Although full and empty capsids were not separated using this protocol, good particles with good shape and high purity were obtained.

As TFF and ceramic hydroxyapatite chromatography have already been used in biopharmaceutical manufacturing, both are scalable methods. The scalability can be adjusted according to the equipment, facility capabilities, and sample requirements.

DNase pre-treatment is usually performed during purification to remove dsDNA derived from host cells. This pre-treatment step is not required, thereby promoting time and cost-effectiveness. However, when DNA removal is insufficient during scale-up, a pre-treatment step should be considered.

### Time Considerations

For Basic Protocol 1, the seeding of cells takes 30 min on Day 1, transfection takes 1 h on Day 4, and harvesting is completed in 40 min on Day 9; thus, a total of 9 days are required to carry out this protocol. For Basic Protocol 2, the preparation (pre-warming buffer and sample, and setting and washing TFF) requires approximately 1 h, the TFF run takes approximately 2–3 h, and the typical cleaning process lasts 1–1.5 h.

For Basic Protocol 3, packing ceramic hydroxyapatite powder into the column takes approximately 5 min and setting up the chromatography system takes 20 min. An additional 30–40 min is required to load the column onto the chromatography system and then wash and equilibrate the column. Purification requires approximately 40 min, and sterilization and cleaning take approximately 20–30 min.

For Basic Protocol 4, qPCR lasts approximately 2.5 h; SDS-PAGE takes approximately 3–4 h; dsDNA detection takes 30 min; and HCP ELISA takes 2.5–3 h. For TEM analysis, approximately 5 min is required to prepare the sample, approximately 1 h is required to set up the TEM system, and approximately 1.5 h is required for observation.

## ACKNOWLEDGMENTS

We thank Dr. Rumi Miyaoka (Asahi Kasei Medical Co., Ltd.) for providing technical support for TFF, and Dr. Yukihiko Hirai (The Institute of Medical Science, The University of Tokyo) and Dr. Tomohiko Yoshitake (HOYA Technosurgical Corporation) for the helpful discussions. This research was funded by the Japan Agency for Medical Research and Development (AMED) with the grant numbers JP21ae0201001, JP21ae0201005, and JSPS KAKENHI with the grant number 20K06464 and 20H03788.

## DATA AVAILABILITY STATEMENT

The data, tools, and materials that support the protocol are available from the corresponding author upon reasonable request.

## INTERNET RESOURCES

https://www.alacrita.com/hubfs/AAV%20Gene%20Therapy%20Landscape%20-%20Sept%202019%20-%20Alacrita.pdf

